# Low cadmium concentrations alter B and T cell responses in Jamaican fruit bats (*Artibeus jamaicensis*)

**DOI:** 10.64898/2026.03.31.715675

**Authors:** Laura A. Pulscher, Phillida A. Charley, Shijun Zhan, Clara Reasoner, Bradly Burke, Tony Schountz

## Abstract

Bats are exposed to a variety of pollutants, including cadmium (Cd), that can impair immune function and potentially increase viral shedding and burden. Despite this, little is known about the impacts of heavy metals on bats. This study aimed to determine the impacts of Cd exposure on bat T and B cell immune responses in naïve and coronavirus infected bats and determine the impact of Cd on viral replication in Jamaican fruit bat (JFB; *Artibeus jamaicensis*) cells. To determine the impact of Cd exposure on adaptive immune responses, splenocyte cultures from naïve and BANAL-52 coronavirus infected JFB were treated with 0, 1, and 10 µM Cd and stimulated overnight with concanavalin A. RNA was extracted, a SYBR Green qPCR was used to assess gene expression. To determine if Cd exposure increased viral replication, two JFB kidney cell clones were treated with 0, 1, 10, and 50 µM of CdCl_2_ overnight and then infected with Cedar virus (CedV). Supernatants were collected and viral titers determined. Several transcripts were upregulated in both naïve and virus infected JFB splenocytes treated with Cd. B cell transcripts were significantly upregulated in a dose-dependent manner and T cell transcripts were also increased in Cd treated splenocytes. Assessment of transcripts associated with T cell subsets suggest a predominant Th2 response in Cd treated splenocytes. Viral replication was not significantly different in Cd treated kidney clones compared to the non-treated cells. These studies provide evidence that JFB adaptive immune responses are altered when exposed to low Cd concentrations.

## 1. Introduction

Bats (order Chiroptera) are reservoir hosts of many viral pathogens of human and animal health importance. While a few viruses, such as rabies virus and other lyssaviruses prove fatal, most infections remain subclinical in bat hosts with seemingly rare shedding. The mechanisms by which bat immune responses control these infections remain poorly understood but likely involves innate mechanisms including dampened inflammasome activity and strong constitutive antiviral immunity, particularly type I interferon responses (1–3). Ecological stressors may compromise immune function and increase viral shedding and burden. For example, climate induced nutritional stress and dietary alterations have been proposed as drivers of Hendra virus spillover (4). Similarly, exposure to non-essential heavy metals, such as cadmium (Cd), could impair immune function, and increase viral replication and shedding in bat hosts, though this remains understudied.

Cadmium is a non-essential heavy metal naturally present in low environmental concentrations. Human activities including mining, manufacturing, and phosphate fertilizer use increase environmental Cd, with consequential health impacts due to its carcinogenic properties and multi-organ toxicity (5). Recent research in rodents demonstrates Cd immunotoxicity through accumulation in immune cells, inducing apoptosis, and altering cytokine expression that may impact pathogen burden and transmission (6). Bats encounter a variety of pollutants, including Cd, due to their wide distributions and diverse dietary habits (7–11). A meta-analysis of tissue metal concentrations in bats found frugivorous bats had significantly increased renal tissue Cd concentrations compared to insectivorous species (10), suggesting increased risk of Cd bioaccumulation compared to non-frugivorous bats. Increased Cd bioaccumulation could impair immunity, increase pathogen burden, and shedding; however, little hypothesis driven work has been performed to determine this.

Few studies have assessed Cd impacts on bat immunity or pathogen burden. Experimental exposure of wild-caught great fruit-eating bats (*Artibeus lituratus*) to low concentrations of Cd (1.5 mg/kg Cd) caused significant liver and kidney dysfunction, kidney leukocyte infiltration, and increased oxidative stress (12). However, prior metal exposure history in wild-caught bats confounded interpretation of these acute responses. Other observational studies of bats captured in polluted sites report elevated heavy metal tissue concentrations, including Cd, were associated with hepatotoxicity and renal toxicity as demonstrated by histopathology, decreased metabolic activity, and increased DNA damage and antioxidant capacity (13–15), all of which could impact bat immunity and pathogen burden. Only one study has examined the impact of Cd on pathogen prevalence and reported lesser horseshoe bats (*Rhinolophus hipposideros*) with increased fecal Cd concentrations correlated positively with *Eimeria hessei* parasite burden (16). In humans, epidemiological studies have identified a link between Cd exposure and increased susceptibility and severity of viral respiratory diseases (17–21). Additionally, influenza A virus replication increased by 1.4 to 2.6 times in a dose-dependent manner in Cd-treated canine kidney cells compared to untreated cells, suggesting Cd exposure may increase pathogen burden (22). As such, the literature suggests immune function and subsequent viral burden, transmission dynamics, and clinical signs of disease could be similarly impacted in Cd exposed bats.

Kidney tropic Cd bioaccumulation in conjunction with kidney tropic bat-borne viruses may have detrimental impacts on viral burden, shedding, and transmission. Henipaviruses such as Hendra, Nipah and Cedar viruses, are negative-sense RNA viruses in the *Paramyxoviridae* family that are naturally hosted by pteropus bats, and have kidney tropism and are secreted through urine (23–26). Hendra and Nipah viruses are high consequence zoonotic pathogens, posing significant public health threats, with high case fatality rates in humans. Cedar virus (CedV), on the other hand, is not known to be pathogenic to humans but shares significant genomic features with Hendra and Nipah virus (24), and can be studied in lower containment – making it a good proxy virus to study viral transmission dynamics of henipaviruses and how these dynamics shift under Cd exposure.

Bat-borne coronaviruses represent another viral group of interest as positive-sense RNA viruses. Interest in bat-borne zoonotic coronaviruses, particularly those in the subgenera *Merbecovirus* and *Sarbecovirus*, has intensified following the emergence of Middle East respiratory syndrome coronavirus (MERS-CoV), severe acute coronavirus 1 (SARS-CoV-1), and SARS-CoV-2 (27). Subsequently, several new bat-borne coronaviruses, including multiple SARS-related coronaviruses (SARSr-CoV), have been discovered and horseshoe bats (*Rhinolophus spp*.) appear to be the natural reservoir for these SARSr-CoVs (28,29). Despite zoonotic coronavirus emergence, coronavirus transmission dynamics within bat populations and potential ecological drivers remain poorly understood. Unlike human-adapted respiratory coronaviruses, bat-borne coronaviruses are primarily enteric viruses, replicating in intestinal tissues and shedding through feces in natural bat hosts (29,30). Considering the gastrointestinal tract is also a primary target of Cd toxicity, causing acute inflammatory responses in the intestines (31), Cd exposure among bats could plausibly increase bat susceptibility to or shedding of enteric pathogens such as coronaviruses, though this remains unstudied.

Nearly all studies of Cd exposure on immune function in bats are observational, apart from Destro et al. (12) which was semi-experimental. Multiple stressors, including environmental pollutants, potentially impact bat immune function and viral replication, making it difficult to isolate Cd effects without controlled studies. Given that bat immune responses differ from laboratory rodents used in most toxicology studies, bats may exhibit different sensitivity or responses to Cd exposure warranting controlled studies in bat models. The Jamaican fruit bat (*Artibeus jamaicensis*) serves as a well-established model for immunology and infectious disease research (32–36), including transmission dynamics of bat-borne pathogens and stress responses (37). Research has developed both *in vitro* and *in vivo* models to examine bat viral responses (38), including coronaviruses (32,36,39), influenza viruses (34,37,40), and paramyxoviruses (41), making this species ideal for studying Cd effects on immune function and viral dynamics in frugivorous bats.

This study aimed to determine Cd impacts on B and T cell immune function and viral replication in frugivorous bats using Jamaican fruit bats as a model. We hypothesized Cd would negatively impact B and T cell function and increase viral replication in a dose-dependent manner. To test this, we treated naïve Jamaican fruit bat splenocyte cultures with various Cd concentrations and conducted a qPCR array targeting various B and T cell gene transcripts. We then opportunistically treated splenocyte cultures from bats infected with BANAL-52 coronavirus (B52-CoV), a SARS-CoV-2 related bat coronavirus, to determine if B and T cells from infected bats would respond similarly to Cd exposure. Finally, we tested Cd effects on viral replication by treating Jamaican fruit bat kidney cell clones with Cd, followed with CedV inoculation.

### 2. Methodology

### 2.1 Jamaican fruit bat kidney cell Cd cytotoxicity assay

To determine the doses used for Cd studies, the cytotoxicity of cadmium chloride (CdCl_2_, Sigma-Aldrich, St. Louis, MO, USA) was first evaluated by a neutral red uptake assay. Briefly, a Jamaican fruit bat polyclonal epithelial cell line (Ajk6) was cultured in a 96 well plate in Dulbecco’s modified Eagle’s medium (DMEM, Gibco), supplemented with 10% fetal bovine serum (FBS, Gibco), penicillin 100 U/mL and streptomycin 100 mg/mL (Gibco). Cells were then treated with 2-fold dilutions of CdCl_2_ (0 to 2,000 µM) for 18, 24 and 48 hours. Cells were then stained with neutral red for 2 hours, washed and then destained. Optical densities were read at 540 nm (neutral red absorbance) and 690 nm (background absorbance). Samples were normalized against the plate background absorbance and percent viability was calculated by taking the difference of the absorbance of the treated cells from the nontreated cells and dividing by the absorbance of the nontreated cells. Percent cytotoxicity was then calculated by subtracting the percent viability from one hundred.

### 2.2 Virus Production and Titration

A molecular clone of B52-CoV was used for this study (42) and was passaged once in Vero E6-TMPRSS2-T2A-ACE2 cells to generate a stock. Briefly, Vero E6-TMPRSS2-T2A-ACE2 cells were cultured in DMEM supplemented with 1% sodium pyruvate, 10% fetal bovine serum (FBS), penicillin 100 U/mL and streptomycin 100 mg/mL at 37°C and 5% CO_2_ and then infected with B52-CoV at a multiplicity of infection (MOI) of 0.1 supplemented with 2% FBS. After infection, the supernatant was collected and stored at -80°C. A molecular clone of a fluorescently labelled Cedar virus (CedV-GFP) was also used for this study (43) and a similar process was used to generate a stock except CedV-GFP was passaged once on Vero E6 cells. Viral titers were measured by TCID_50_ using Vero E6-TMPRSS2-T2A-ACE2 cells for B52-CoV and Vero E6 cells for CedV-GFP.

### 2.3 Jamaican fruit bats

Eight male Jamaican fruit bats were selected for the first part of the study assessing the impact of Cd on immune function. These bats were selected from our Jamaican fruit bat breeding colony which is maintained in a free-flight facility. After collection, bats were euthanized, and spleens were collected to make single cell suspensions. Additionally, we opportunistically collected spleen from 3 bats who were inoculated with B52-CoV for another study. Bats were inoculated with 10^5^ TCID_50_ of virus while holding the bats in a contralateral position and intranasally inoculated with 10^5^ TCID_50_ equivalents per bat in 50 µL 2% FBS DMEM. Infected bats were confined in bird cages under BSL-3 containment. On day 12, bats were euthanized, and spleens collected to make single cell suspensions. This study was performed with approval from the Colorado State University Institutional Animal Care and Use Committee.

### 2.3 Jamaican fruit bat splenocyte culture preparation and cadmium treatment

For the single cell splenocyte suspensions, spleens were collected and immediately placed on ice with Clicks medium (Gibco). Splenocytes were cultured by gently disrupting the spleen in a 100 µm cell strainer and washing cells through the strainer with Clicks medium (Fujifilm). Red blood cells were then lysed with ammonium chloride buffer, followed by two washes with Clicks medium. Splenocytes were then counted, and six identical cultures were plated for each bat with 2.5 x 10^5^ to 5.0 x 10^6^ splenocytes each, apart from one of the B52-CoV inoculated bats (B52-CoV Bat 4) which only had enough splenocytes for three identical replicates. Apart from the single B52-CoV bat with one replicate for each treatment, duplicate cultures from each bat were treated with concentrations of 0, 1 or 10 µM CdCl_2_ and 2.5 µg of concanavalin A for 15 hours in 10% FBS Clicks medium supplemented with 10% FBS at 37°C and5% CO_2_. Splenocyte cultures were collected and total RNA was extracted with the Qiagen RNeasy Mini Kit (Qiagen) and frozen at -80°C until further processed.

### 2.4 Immune gene expression profiling

Total RNA was extracted from splenocytes with the Qiagen RNeasy Mini Kit and reverse transcribed into cDNA (QuantiTech RT, Qiagen). Gene expression was determined using SYBR Green qPCR (QuantiTech SYBR Green, Qiagen) arrays as previously described (36). Primers for Jamaican fruit bat genes are described in Supplementary Table 1. Within sample gene normalization was performed on Rps18 (ΔCq), and fold-change determined by comparing each gene from the Cd treated cells to the non-Cd treated cells from the same bat (ΔΔCq). A cycle threshold (C_t_) cut off value of 35 was used for all genes. Fold-change values were assessed for normality and log 2 transformed. In some instances, RNA was degraded so a replicate had to be thrown out of the dataset.

### 2.5 Cedar virus replication in CdCl_2_ treated Jamaican fruit bat kidney cells

To determine the impact of Cd on viral replication in bat cells; Jamaican fruit bat kidney cell clones, Ajk6-2 and Ajk6-10, were cultured in quadruplicate in DMEM, supplemented with 10% FBS, penicillin 100 U/mL and streptomycin 100 mg/mL. Ajk6 cell clones are primary kidney epithelial cells that were generated from a single Jamaican fruit bat kidney and then single cell sorted to expand individual clones (unpublished data). Ajk6 cell clones were then treated overnight with 0, 1, 10, or 50 µM of CdCl_2_ and two replicates of each of the Ajk6 cell clones were infected with a 0.1 multiplicity of infection (MOI) of CedV-GFP. The other replicates were mock infected with viral media. Infected and mock-infected cells were incubated for 30 min at 37°C with rocking every 10 min, washed with PBS and then incubated with DMEM supplemented with 2% FBS, 1.5% HEPES, and 100 U/mL penicillin and 100 mg/mL streptomycin. Photos were taken and supernatant was collected at 1 hour post infection (hpi) and every 24 hours for 3 days. Viral titers of the supernatants were determined using a TCID_50_ assay. This experiment was repeated three separate times for each clone.

### 2.6 Data Analysis

To determine a toxic dose for Cd in Ajk6 cells, a TD_50_ was determined by a neutral red uptake assay and calculated by fitting cadmium response data to a four-parameter logistic equation. For the gene expression analysis from splenocytes cultured from naïve bats, four replicates were removed from the analysis due to degraded RNA including one replicate for bat 3 for the 0 µM CdCl_2_ treatment group, one replicate for bat 6 for the 1 µM CdCl_2_ and 10 µM CdCl_2_ treatment group, and one replicate for bat 8 for the 10 µM CdCl_2_ treatment group. Additionally, one replicate from B52-CoV infected bat 3 was removed for the 10 µM CdCl_2_ treated group due to degraded RNA. Therefore, the data from a single replicate was used for each of these treatments for these bats in subsequent analysis. To determine differences in immune gene expression profiling between the 0, 1, and 10 µM CdCl_2_ treated groups, a heat map was first generated to assess overall differences across treatment group and gene expression and to visually detect outliers. A mixed-effects analysis with Geisser-Greenhouse correction and a Tukey’s multiple comparison test was used to determine statistical significance between the different treatment groups for each gene transcript. A repeated-measures one-way ANOVA with Geisser-Greenhouse correction and Tukey’s multiple comparison test was used to determine statistical significance in CedV titers in Ajk6-2 and Ajk6-10 cell clones treated with 0, 1, 10, and 50 µM of CdCl_2_ at each timepoint (0, 24, 48, and 72 hpi). All statistical analyses were performed using GraphPad Prism version 10.4.1 for Windows (GraphPad Software, Boston, Massachusetts USA, www.graphpad.com) or R version 2026.01.1 for Windows. Statistical significance was considered at α ≤ 0.05.

## 3. Results

### 3.1 CdCl_2_ concentrations ≥32 µM is cytotoxic to Jamaican fruit bat kidney cells

Treatment with ≤ 32 µM of CdCl_2_ was not cytotoxic to Ajk6 cells after 18, 24 or 48 hours of exposure (Figure 1; Supplementary Figure 1). Cadmium was cytotoxic to Ajk6 cells at a CdCl_2_ concentration of ≥ 250 µM after 18 and 24 hours of exposure. A TD_50_ of 364.7 µM of CdCl_2_ and 368.5 µM of CdCl_2_ was calculated for cells treated for 18 and 24 hours, respectively. Concentrations of ≥ 62 µM of CdCl_2_ were cytotoxic to cells after 48 hours of exposure with a TD_50_ of 139 µM of CdCl_2_.

**Figure 1.**
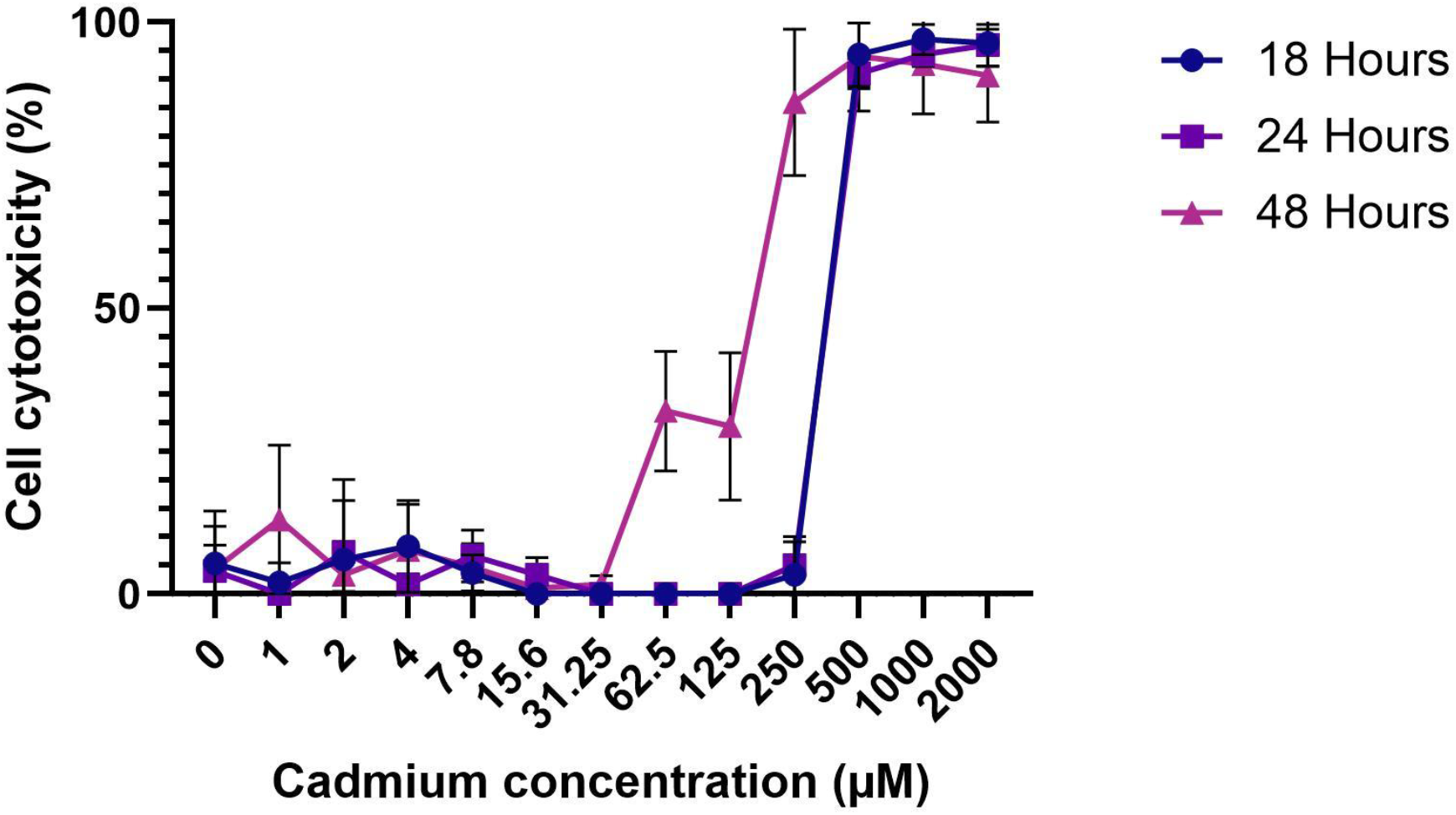
Percent (%) cytotoxicity of Ajk6 polyclonal cells treated with 0 - 2,000 µM of CdCl_2_ over 18 (blue line with circles), 24 (purple line with squares) or 48 (pink line with triangles) hours.

### 3.2 B, Th1 and Th2 cells are altered in a dose-dependent manner by CdCl_2_

To determine the impact of Cd on T and B cells, replicate splenocyte cultures were treated with 0, 1, or 10 µM CdCl_2_, stimulated with concanavalin A, and RNA extracted for RT-qPCR for B and T cell gene expression profiling. Bat splenocyte cultures treated with CdCl_2_ had increased expression of several genes compared to non-Cd treated cultures apart from bat 6 which had decreased gene expression for all genes except for T cell receptor β (*Tcrβ*) suggesting this bat was an outlier (Figure 2 and 3). Subsequent analyses included gene expression analysis of bats 1 to 5, 7, and 8.

**Figure 2.**
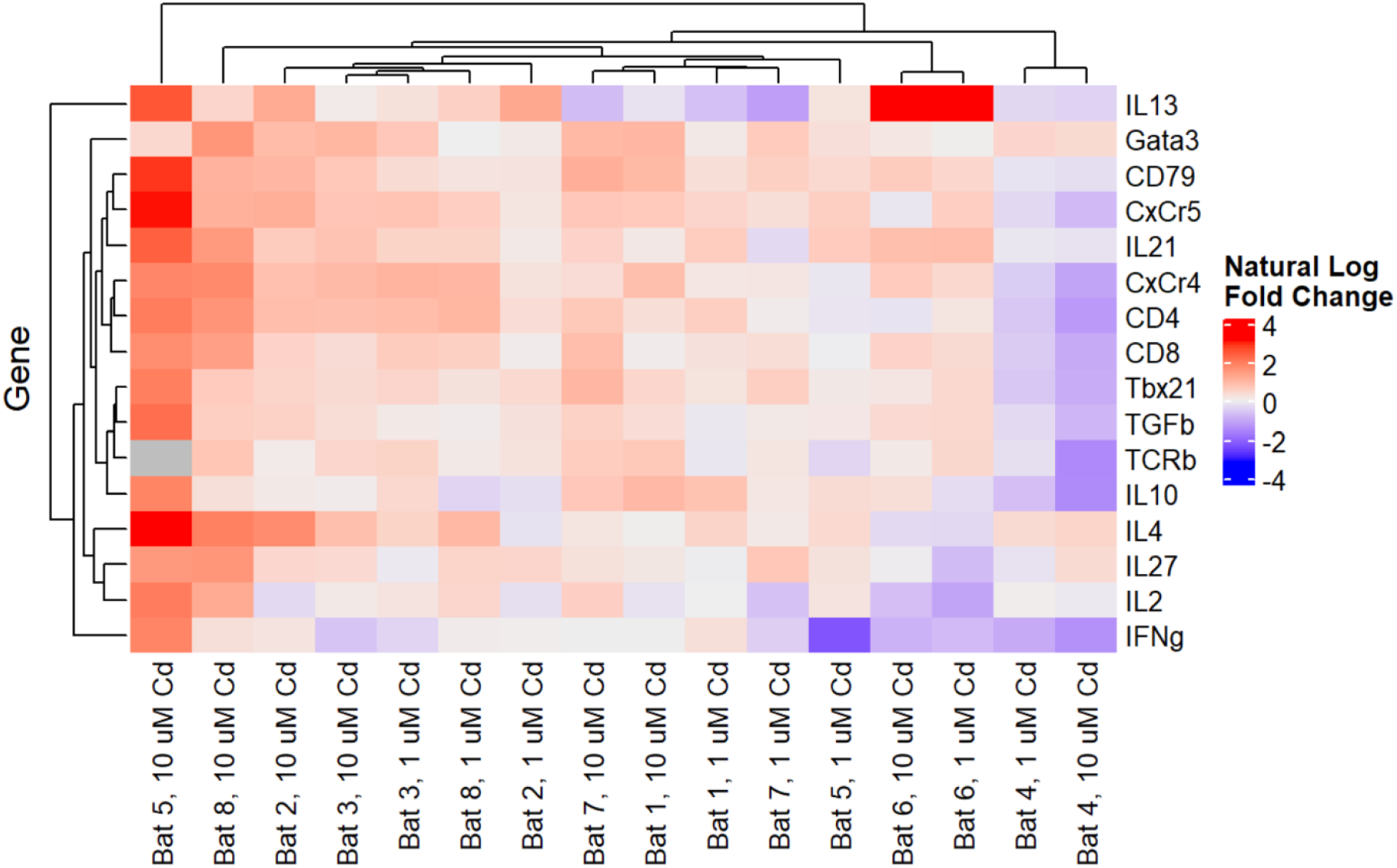
Heatmap of gene expression for Cd-treated Jamaican fruit bat splenocytes treated with 1 µM or 10 µM of CdCl_2_. Fold-change was determined by comparing each transcript from the Cd treated cells to the non-Cd treated cells from the same bat.

**Figure 3.**
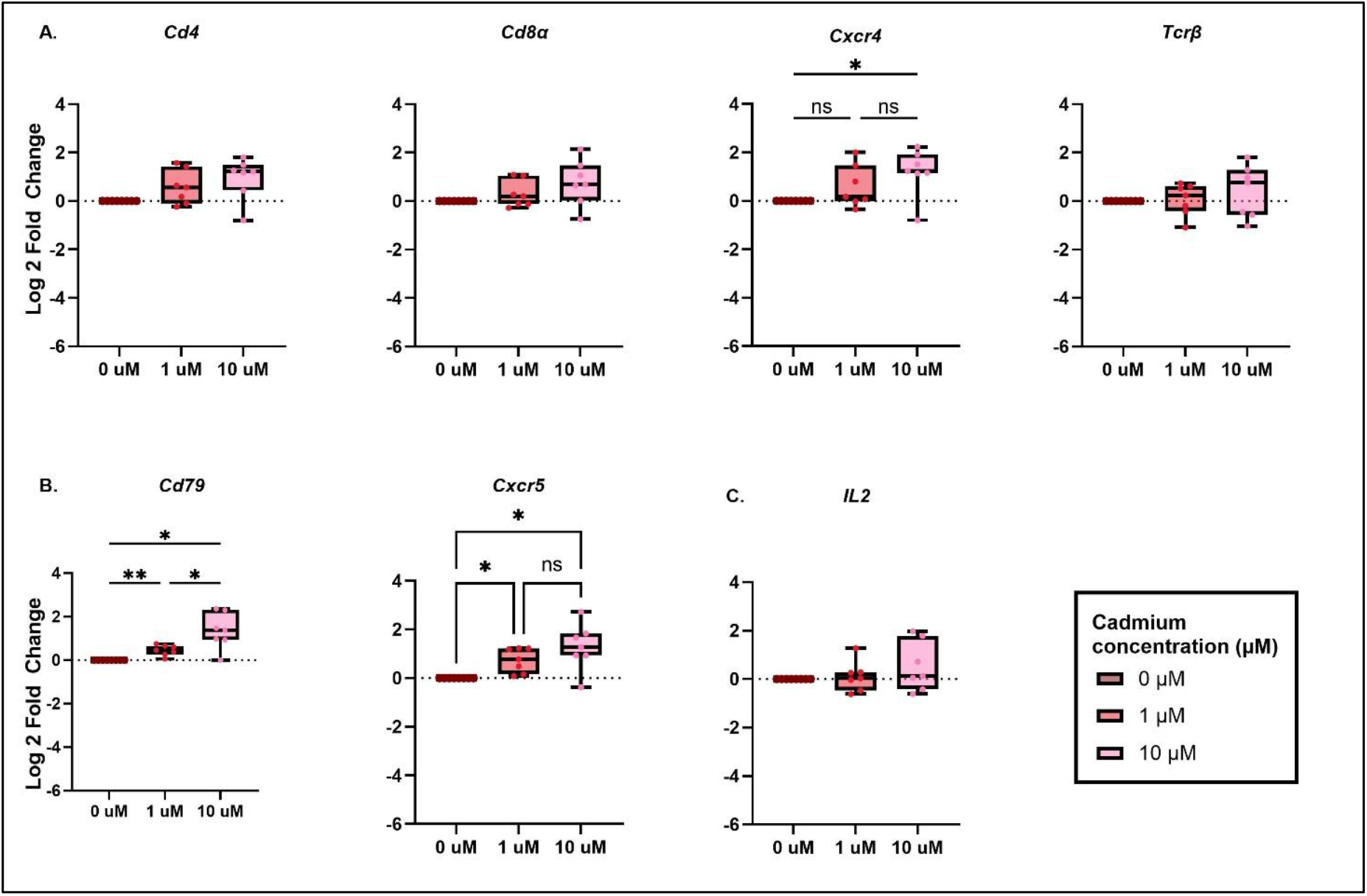
Differential expression for Cd-treated Jamaican fruit bat splenocytes for A) T cells, B) B cells, and C) *IL2* cytokine gene transcripts for splenocytes treated with 0 µM (dark pink), 1 µM (salmon), and 10 µM (light pink) CdCl_2_. The dashed line represents baseline expression for non-Cd treated cells (0 µM CdCl_2_ treated group). Fold-change was determined by comparing each transcript from the Cd treated cells to the non-Cd treated cells from the same bat. A mixed-effects analysis with the Geisser-Greenhouse correction and Tukey’s multiple comparisons test was used to evaluate significant differences in gene expression between cadmium treatment groups. *p-value ≤ 0.05, **p-value ≤ 0.01, and ns = not significant.

Gene expression was increased, or trending towards increasing in a dose-dependent manner, for B and T cell transcripts (Figure 3). *Cd4* transcripts were significantly higher in Cd-treatment groups for *Cd4* (ε = 0.94, F = 5.6, *p* = 0.02) and approaching significance for *Cd8α* (ε = 0.76, F = 3.7, *p* = 0.08). However, *post hoc* tests were not significantly different between the groups but *Cd4* gene expression between 0 µM CdCl_2_ and 10 µM CdCl_2_ treatment groups were approaching significance (mean difference = -0.90, SE = 0.33, *p* = 0.06, 95% CI[-1.95,0.07]). *Cxcr4* gene expression was also significantly increased in the 10 µM CdCl_2_ group compared to the non-Cd treated cells (mean difference = -1.20, SE = 0.37, *p* = 0.04, 95% CI[-2.33,-0.07]). Gene transcripts for *Tcrβ* (ε = 0.64, F = 0.66, *p* = 0.47) and *IL2* (ε = 0.75, F = 1.66, *p* = 0.24) were not significantly different across the different treatment groups. B cell *Cd79* gene expression was significantly increased in both Cd treated groups compared to the non-Cd treated group (0 vs 1 µM CdCl_2_ group: mean difference = -0.45, SE = 0.09, *p* = 0.01, 95% CI[-0.73,-0.17]; 0 vs 10 µM CdCl_2_ group: mean difference = -1.35, SE = 0.31, *p* = 0.01, 95% CI[-2.30,-0.39]). There was also a significant dose-dependent increase in *Cd79* expression between the 1 and 10 µM CdCl_2_ group (mean difference = -0.89, SE = 0.28, *p*= 0.04, 95% CI[-1.76,-0.03]). There was also significantly increased gene expression for *Cxcr5* between the non-Cd treated cells and the Cd treated cells (0 vs 1 µM CdCl_2_ group: mean difference = -0.73, SE = 0.19, *p* = 0.02, 95% CI[-1.32,-0.15]; 0 vs 10 µM CdCl_2_ group: mean difference = -1.29, SE = 0.36, *p* = 0.03, 95% CI[-2.40,-0.17]).

To determine the specific T helper (Th) cell response, we also examined expression of transcripts involved in Th1, Th2, and T regulatory (Treg) cell responses. Transcripts associated with a Th1 response were significantly different between treatment groups for *IL27* (ε = 0.58, F = 8.97, *p* = 0.02) and *Tbx21* (ε = 0.76, F = 5.34, *p* = 0.04). *Post hoc* analysis found *IL27* expression remained significantly increased between the Cd treated and the non-Cd treated groups (0 vs 1 µM CdCl_2_ group: mean difference = -0.33, SE = 0.10, *p* = 0.03, 95% CI[-0.64,-0.03]; 0 vs 10 µM CdCl_2_ group: mean difference = -1.00, SE = 0.31, *p* = 0.04, 95% CI[-1.94,-0.05]). While not significantly different, *IL27* expression was increasing in a dose-dependent manner between the Cd treated groups and was approaching significance (1 vs 10 µM CdCl_2_ group: mean difference = -0.66, SE = 0.29, *p* = 0.14, 95% CI[-1.55,0.23]). *Tbx21* gene expression was not significantly different between the treatment groups for the *post hoc* analysis but the Cd-treated groups were trending towards increased expression compared to the non-Cd treated group and were approaching significance (0 vs 1 µM CdCl_2_ group: mean difference = -0.46, SE = 0.19, *p* = 0.12, 95% CI[-1.05,0.14]; 0 vs 10 µM CdCl_2_ group: mean difference = -0.80, SE = 0.32, *p* = 0.10, 95% CI[-1.09,0.17]).

For the Th2 gene transcripts, gene expression was significantly different between the treatment groups for *Gata3* (ε = 0.56, F = 14.75, *p* = 0.003), *IL4* (ε = 0.67, F = 8.74, *p* = 0.01), and *IL21* (ε = 0.60, F = 8.61, *p* = 0.02) but not *IL13* (ε = 0.54, F = 0.13, *p* = 0.75). *Post hoc* analysis found *Gata3* expression was significantly increased in the Cd treated groups compared to the non-Cd treated group (0 vs 1 µM CdCl_2_ group: mean difference = -0.59, SE = 0.09, *p* = 0.002, 95% CI[-0.87,-0.31]; 0 vs 10 µM CdCl_2_ group: mean difference = -1.23, SE = 0.30, *p* = 0.02, 95% CI[-2.22, 0.24]) but *Gata3* expression was not significantly different between the 1 and 10 µM groups (mean difference = -0.64, SE = 0.32, *p* = 0.20, 95% CI[-1.67,0.40]). Similarly, *IL21* expression was significantly increased in Cd treated groups compared to the non-Cd treated control (0 vs 1 µM CdCl_2_ group: mean difference = -0.47, SE = 0.15, *p* = 0.05, 95% CI[-0.92,-0.01]; 0 vs 10 µM CdCl_2_ group: mean difference = -1.28, SE = 0.41, *p* = 0.04, 95% CI[-2.53, -0.04]); however, *IL21* expression was not significantly different between the 1 and 10 µM CdCl_2_ groups (mean difference = -0.82, SE = 0.37, *p* = 0.15, 95% CI[-1.95, 0.31]). Gene expression for *IL4* was not significantly different between the 0 and 1 µM CdCl_2_ group (mean difference = -0.48, SE = 0.33, *p* = 0.38, 95% CI[-1.50, 0.54]) but *IL4* was significantly increased in the 10 µM CdCl_2_ treated group compared to the 0 and 1 µM CdCl_2_ groups (0 vs 10 µM CdCl_2_ group: mean difference = -1.41, SE = 0.46, *p* = 0.05, 95% CI[-2.81, - 0.01]; 1 vs 10 µM CdCl_2_ group: mean difference = -0.93, SE = 0.24, *p* = 0.02, 95% CI[-1.67, -0.20]).

The Treg associated gene transcripts did not significantly differ between the Cd treatment groups for *Foxp3* (ε = 0.58, F = 0.46, *p* = 0.54), *IL10* (ε = 0.73, F = 0.53, *p* = 0.55) or *Tgfβ* (ε = 0.55, F = 0.29, *p* = 0.63).

### 3.3 Jamaican fruit bats infected with B52-CoV had increased splenic B cell gene transcripts upon exposure to CdCl_2_

To determine the impact of CdCl_2_ on immune function in bats that are infected with a virus, we opportunistically collected and cultured splenocytes from 3 bats inoculated with B52-CoV and then exposed these to CdCl_2_ and ran a B and T cell gene expression array. While not significant, *Cd4* and *Cd8α* gene transcripts in Cd treated splenocytes appeared to have similar or decreased expression compared to non-Cd treated cells (Supplementary Figure 2). However, *Cxcr4* and *Tcrβ* appeared to have increased expression in Cd-treated cells compared to non-Cd treated cells and *Cxcr4* expression was approaching significance (ε = 0.90, F = 2.38, *p* = 0.16). B cell genes had increased expression in Cd treated cells compared to non-Cd treated cells and were approaching significance for both *Cd79* (ε = 0.51, F = 1.44, *p* = 0.09) and *Cxcr5* (ε = 0.59, F = 3.67, *p* = 0.18) (Supplementary Figure 1). While not significant, B52-CoV infected splenocytes treated with CdCl_2_ appeared to have a predominant Th2 response as opposed to a Th1 or Treg response (Supplementary Figure 3).

### 3.4 Treatment of Jamaican fruit bat kidney cells with CdCl_2_ did not impact CedV replication

To determine if low CdCl_2_ concentrations would impact viral replication in bat cells, two Jamaican fruit bat kidney cell clones were treated with 0, 1, 10, or 50 µM of CdCl_2_ and then inoculated with a 0.1 MOI of CedV. A repeated-measures ANOVA revealed there was not a significant difference in CedV titer between the different Cd treatment groups for each time point for either kidney cell clone (Figure 4).

**Figure 4.**
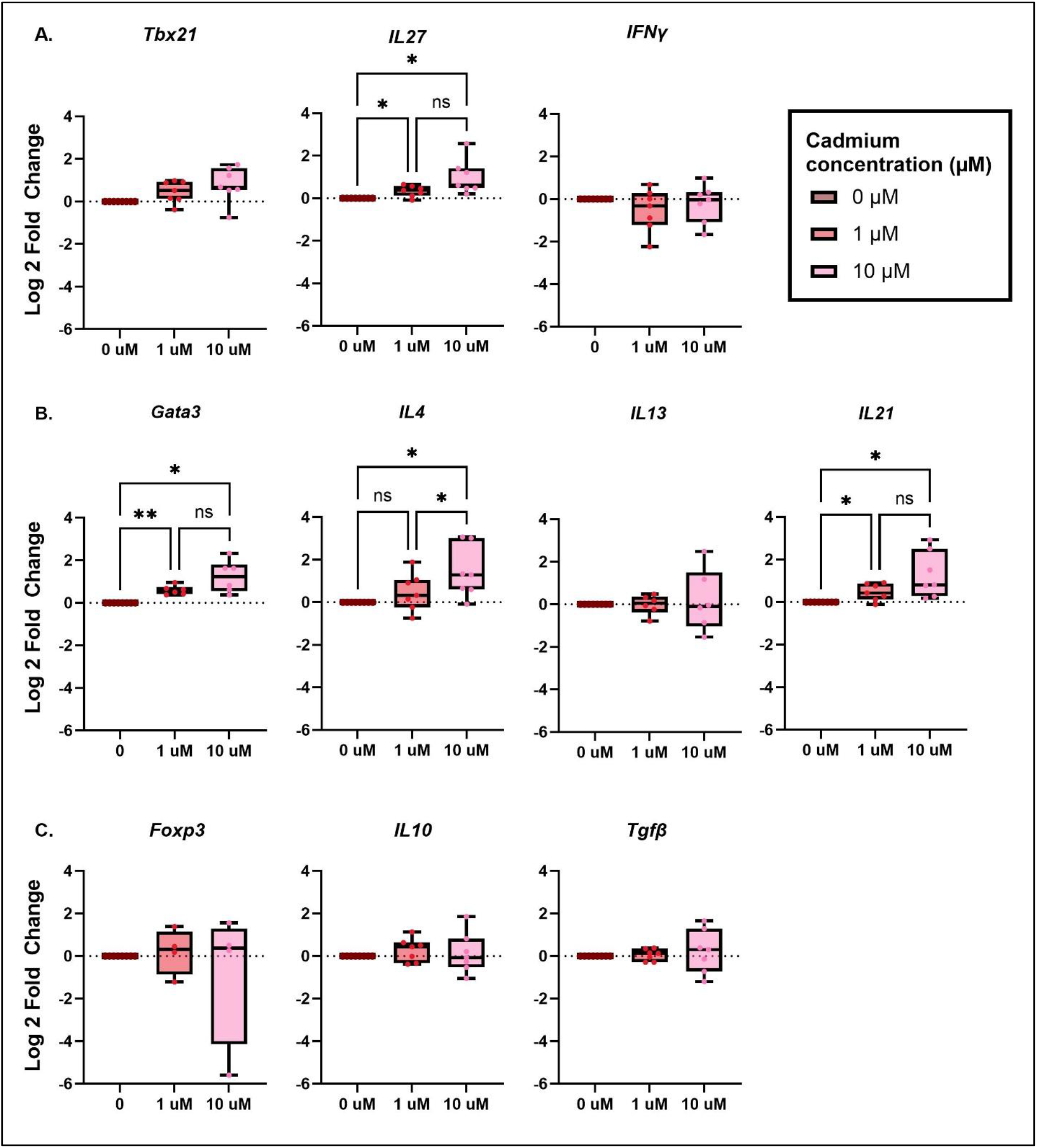
Expression in Cd-treated Jamaican fruit bat splenocytes for A) T helper 1 (Th1), B) Th2, and C) regulatory T cell (Treg) gene transcripts. The dashed line represents baseline expression for non-Cd treated cells (0 µM CdCl_2_ treated group). Fold-change was determined by comparing each transcript from the Cd treated cells to the non-Cd treated cells from the same bat. A mixed-effects analysis with the Geisser-Greenhouse correction and Tukey’s multiple comparisons test was used to evaluate significant differences in gene expression between cadmium treatment groups. *p-value ≤ 0.05 and ns = not significant.

**Figure 5.**
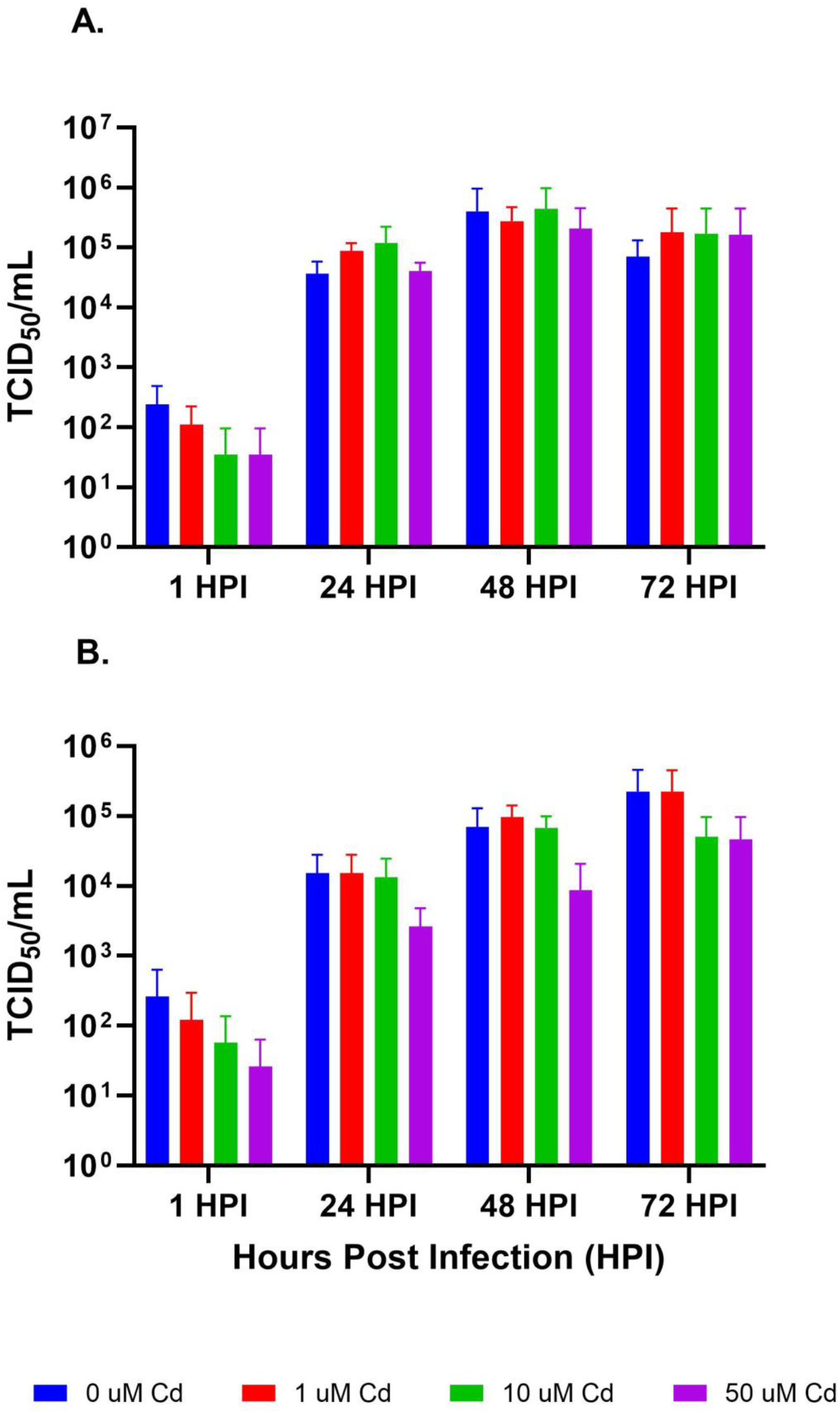
Cedar virus (CedV) replication (TCID_50_/mL equivalent) in Ajk6-2 (panel A) and Ajk6-10 (panel B) cell clones treated with 0 (blue line), 1 (orange line), 10 (gray line), or 50 (yellow line) µM of CdCl_2._ A mixed-effects analysis with the Geisser-Greenhouse correction and Tukey’s multiple comparisons test was used to evaluate significant differences in gene expression between cadmium treatment groups at each time point.

## 4. Discussion

Recent studies have identified increased concentrations of heavy metal contaminates, including Cd, among bat species that may impact their overall health, immune function, and subsequently viral transmission dynamics. This study sought to determine the impact of Cd on B and T cell immune responses and viral replication in Jamaican fruit bats. Our results suggest that even low levels of Cd alter Jamaican fruit bat B and T cell responses in a dose-dependent manner. In particular, B cell gene transcripts had increased expression in Cd-treated fruit bat splenocytes compared to non-Cd treated splenocytes. This is supported by the significantly increased expression of *Cd79* and *Cxcr5*, both of which are highly expressed on B cells, in both Cd-treated groups compared to the non-Cd treated splenocytes as well as a significantly increased expression of *Cd79* in the 10 µM Cd treated splenocytes compared to the 1 µM Cd treated splenocytes, suggesting a dose-dependent response. This is opposite of what previous *in vitro* studies report where Cd exposure was reported to induce B cell apoptosis in a dose-dependent manner (6). However, *in vivo* studies report mixed results with low concentrations of Cd reducing the number of B cells and high Cd concentrations increasing B cell numbers (6). One study also reported Cd concentrations <1 mg Cd/kg led to a decrease of blood B cells but an increase in splenic B cells (44) which might explain why we found increased expression of gene transcripts associated with splenic B cells in this study.

While not significantly different, *Cd4* and *Cd8α* transcripts were increased in a dose-dependent manner in Cd-treated splenocytes compared to non-Cd treated splenocytes, with increased expression of *Cd4* cells compared to *Cd8α* cells. There was also 2-fold and 1.4-fold increased expression of *Cd4* and *Cd8α*, respectively, in splenocytes treated with 10 µM Cd compared to those treated with 1 µM Cd. The increased expression of *Cd4* and *Cd8α* is also supported by the increased expression of *Cxcr4* and *Tcrβ*, both of which are expressed by T cells, in a dose-dependent manner with cells treated with 10 µM Cd having a significantly higher expression of *Cxcr4* compared to cells treated with 0 µM of Cd. Opposite of what would be expected, *IL2*, an important cytokine for T cell proliferation, expression was similar across all treatment types (Cd-treated and non-Cd treated cells). While we are unsure why this is, other cytokines, including IL4 or IL15, can drive T cell proliferation for specific T cell types. It is possible that these other cytokines may be leading to increased T cell proliferation in this case as *IL4* had increased expression in the Cd-treated splenocytes compared to the non-Cd treated splenocytes. Together these data suggest that exposure to low Cd concentrations leads to increased gene expression in splenic T cells. This is opposite of what has been reported in other *in vitro* studies which report overall decreased T cell proliferation in mouse splenocytes (45), or increased apoptosis of CD4 but increases in CD8 which resulted in overall decreases in the CD4/CD8 ratio in a dose- and time-dependent manner in murine thymocytes (46). Another study analyzed the effects of Cd on blood cells and found that CD4 and CD8 cell counts were decreased when exposed to 5 or 10 ppm of Cd but increased at 25 ppm Cd which may be due to increased recruitment of these T cells against Cd toxicants (47). A previous study of acute Cd exposure in wild-caught greater fruit-eating bats suggests that *Artibeus* bats may be more sensitive to heavy metal exposure compared to laboratory rodents (12) which may explain why *Cd4* and *Cd8α* expression was increased in splenocytes treated with low concentrations of Cd in the current study. CD4 and CD8α are also expressed on other cell types such as macrophages and granulocytes (CD4) and natural killer (NK) cells (CD8α) which may also explain the increased expression seen for these gene transcripts in the current study. Future studies using flow cytometry are warranted to determine the proportion of CD4 and CD8α cells as well as the expression levels of these on different cell populations to fully determine the impact of Cd on immune responses in bats.

To further understand the impact of Cd on specific T cell responses, cytokine and chemokine expression for transcripts associated with Th1, Th2, or Treg responses was also determined. Overall, splenocytes treated with low Cd concentrations had increased but mixed Th1 (*Tbx21* and *IL27*) and Th2 (*Gata3, IL4* and *IL21*) responses compared to non-Cd treated splenocytes. Genes associated with a Th2 response were overall increased, albeit slightly, compared to Th1 genes which suggest that a Th2 response is the dominant T cell driving the immune response. Additionally, *IFNƴ*, a cytokine typically thought to drive Th1 responses, had decreased expression in Cd treated cells compared to the non-Cd treated cells and *IL4*, a cytokine typically thought to drive Th2 responses, was significantly increased between the 0 and 10 and the 1 and 10 µM Cd treated groups providing further support that Th2 is the predominant response in this case. These findings are supported by another study of great Himalayan leaf-nosed bats (*Hipposideros armiger*) that found increased expression of *IL6, IL10*, and *TNFα* but decreased expression of *IFNγ* in liver tissues collected from bats in polluted areas in China compared to bats in less polluted areas suggesting an inflammatory response was triggered and Th2 cytokines were released, decreasing cytotoxic T cells (13). Although bats collected in polluted sites had increased Cd liver concentrations, the authors were unable to conclude that the immune responses were driven by Cd exposure alone because increased liver mercury (Hg) and lead (Pb) concentrations were also identified in these bats (13). Regardless, this provides data supporting that heavy metal exposure may drive a predominant Th2 response in bats. While data is mixed, other *in vitro* studies report a similar Th2 response in response to Cd exposure (6,46,48,49) further supporting our findings. The increased expression of *IL4, IL21*, and *Cxcr5* might also indicate that T follicular helper (Tfh) cells are responding, particularly because *CxCr5*, a key receptor on Tfh cells, and *IL21*, a common Tfh cytokine, were also increased in Cd-treated cells compared to the non-Cd treated cells in a dose-dependent manner. Tfh cells are known to work closely with Th2 cells and help guide and support B cell development and antibody formation. Gene expression for Treg transcripts (*Foxp3, IL10*, and *Tgfβ*) did not appear to be significantly altered in the Cd-treated splenocytes compared to the non-Cd treated splenocytes, although there was a lot of variability for *Foxp3* and *Tgfβ* for the splenocytes treated with 10µM Cd. Though the effects of Cd on Treg cells are not well understood, other *in vitro* studies report no effect or a decrease in IL10 production in cells treated with Cd (6,48), which is similar to findings in this study.

To determine the impact of Cd on bats infected with a coronavirus, spleens were opportunistically collected and cultured from bats infected with B52-CoV and then treated with 0, 1 or 10 µM of Cd. While it is difficult to make generalizations on a small data set; overall, similar trends were found in infected bats, with increased expression of B cell associated gene transcripts and a more dominant Th2 response. The Th1 response appeared to be more suppressed in B52-CoV infected bat splenocyte cultures compared to uninfected bats with a lower expression of *IL27*, in both Cd treated groups, and *IFNγ*, in the group treated with 10 µM Cd, compared to non-Cd treated cells.

To better understand the impact of Cd on viral replication in bats, Jamaican fruit bat kidney cell clones were treated with 0, 1, 10 or 50 µM of Cd and then infected with CedV. In this study, CedV replication was not different between the different Cd treatment groups for either of the Ajk6 cell clones. Few studies have determined the impact of Cd exposure on viral replication or shedding among animal species but epidemiological studies among humans have found a correlation between Cd exposure and increased susceptibility and severity of viral respiratory diseases (17–21). Additionally, one *in vivo* study reported Influenza A virus replication was significantly increased in MDCK cells treated with Cd (15 – 50 µM) compared to untreated cells (22). While we are uncertain why an increased CedV titer was not observed in cells treated with lower Cd concentrations in this study, it is possible that the Cd concentrations tested were not high enough to increase viral burden *in vitro*. Alternatively, Cd may not impact the cell processes used by CedV for viral replication. Considering *in vitro* studies do not always align with what is observed *in vivo* and considering Jamaican fruit bat B and T cells exposed to low Cd concentrations are altered *in vitro* it is possible that increased viral burden and shedding would be observed in animal models warranting future *in vivo* studies in Jamaican fruit bats.

## 5. Conclusion

Due to their dietary diversity, bats are exposed to a variety of environmental pollutants, including Cd. This may alter bat immune responses and subsequently lead to increased viral replication or shedding which could ultimately increase opportunities for viral spillovers into human or domestic animal populations. To date, the few studies that have assessed the impact of Cd on bat immune responses and pathogen transmission dynamics have focused on observational studies or toxicological studies of wild-caught bat species, limiting our understanding of the impact of heavy metal pollutants on bats. The current study utilized a Jamaican fruit bat model to provide additional evidence that even at low Cd concentrations, Jamaican fruit bat adaptive immune responses were altered. While this study did not find evidence that acute Cd exposure affected CedV replication, additional *in vivo* studies are warranted to further understand the impact of Cd, and other heavy metals, on immune function and viral transmission dynamics in fruit bats.

## Acknowledgements

We would like to thank Dr. Craig Wilen at Yale University who provided the B52-CoV molecular clone and Dr. Eric Laing at the Uniformed Services University who provided the CedV-GFP molecular clone. We would like to thank Madeline Yunker and Jordyn Walker who assisted with some parts of the laboratory work for this study. The study was supported by internal Colorado State University funds from the College of Veterinary Medicine and Biomedical Sciences.

