## Supplementary Information for "Low cadmium concentrations alter B and T cell responses in Jamaican fruit bats (*Artibeus jamaicensis*)"

**Supplementary Table 1:** Jamaican fruit bat immune gene expression primers, 5' to 3'.

| GENE | FORWARD | REVERSE |
|---|---|---|
| ***RPS18*** | GCGAGTACTCAACACCAATA | TTTTGGTGAGGTCGATGTCT |
| ***IL-2*** | CGAACTCAACCCTCTGAAGG | GTGCGTTCACTGTGACATTG |
| ***IL-4*** | AGCTCTGGTCTGCTTACTAG | TGGTCTCTTCTAAGGTGAGG |
| ***IL-10*** | AGCTCAGCACTTCTCTGTTG | TCTCAGATAGCCTGCTCTGC |
| ***IL-13*** | CATCCAAAAGACCCAGAGG | ATCACTTCCACTTTGGTGTC |
| ***IL-21*** | AGATAATCAGCCTGCTAACG | CTCTGTTTCTGTCTTCTCCC |
| ***IL-27*** | GACCGTGAGTTTGGATCTCC | TCAGGGAGGTTGAATCCTGT |
| ***IFNΓ*** | CCTTGAAAGACAACCAGAGC | CCTCCATTTTGCTGTTGCTG |
| ***CD4*** | CTCATTCCCACTCACCTTCG | CTCCAAGGAGAAGGCAATCC |
| ***CD8A*** | CCCAGATCTCAGGCCAAAAG | AGCCTTGGTCCTTTTCTGTG |
| ***CD79*A** | GGGACAGTCTTCCTCCTCCT | TCCCCCAGACTCACCATCAT |
| ***CXCR4*** | GCTCAGGCGACTATGACTCC | TGCCCACTATGCCAGTCAAG |
| ***CXCR5*** | GGTCAGCCAACCTCATCACA | TAGAGGAAGCGGGAGGTGAA |
| ***TCRB*** | CAACAGCACTGACTCCAACT | CTGTCCAGCCCATAGAACTG |
| ***TGF*B** | TGTTCTTCAACACGTCGGAG | CGTGCTGTTCCACTTTCAAC |
| ***TBX21*** | CCAAAGGATTCCGGGAGAAC | AATTGACAGTTGGGTCCAGG |
| ***GATA3*** | ACCCCTGACTATGAAGAAGG | CACCTTTTTGCACTTTTTGG |
| ***FOXP3*** | CCAGCTCTGTGATCTGGAAC | AAAGCCTGTTGAGGTCCATC |

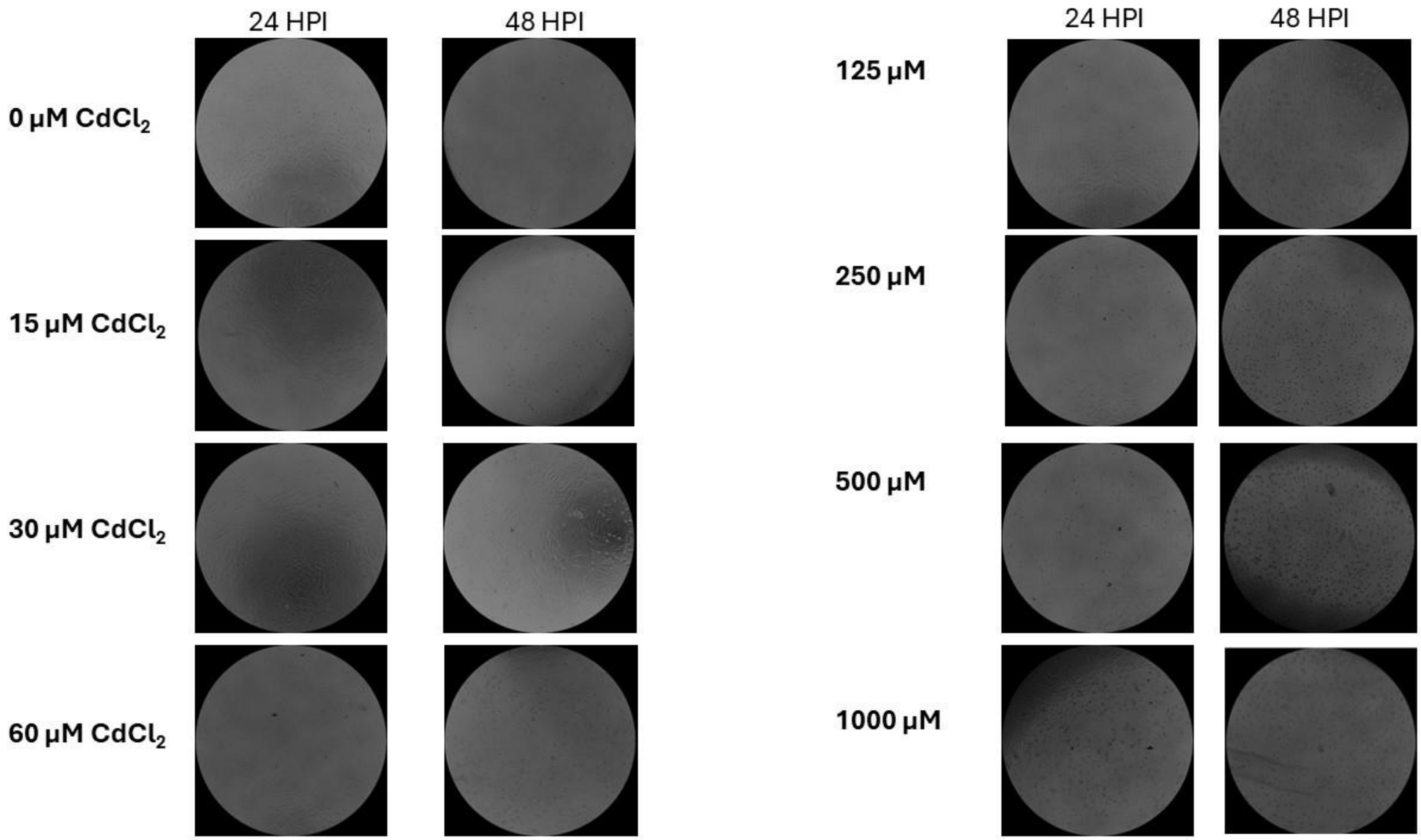


**Supplementary Figure 1:** Microscopic images of *Artibeus jamaicensis* kidney cells (Ajk6) treated with various cadmium concentrations (range of 0 – 1000 µM) for 24 hours (left) or 48 hours (right).

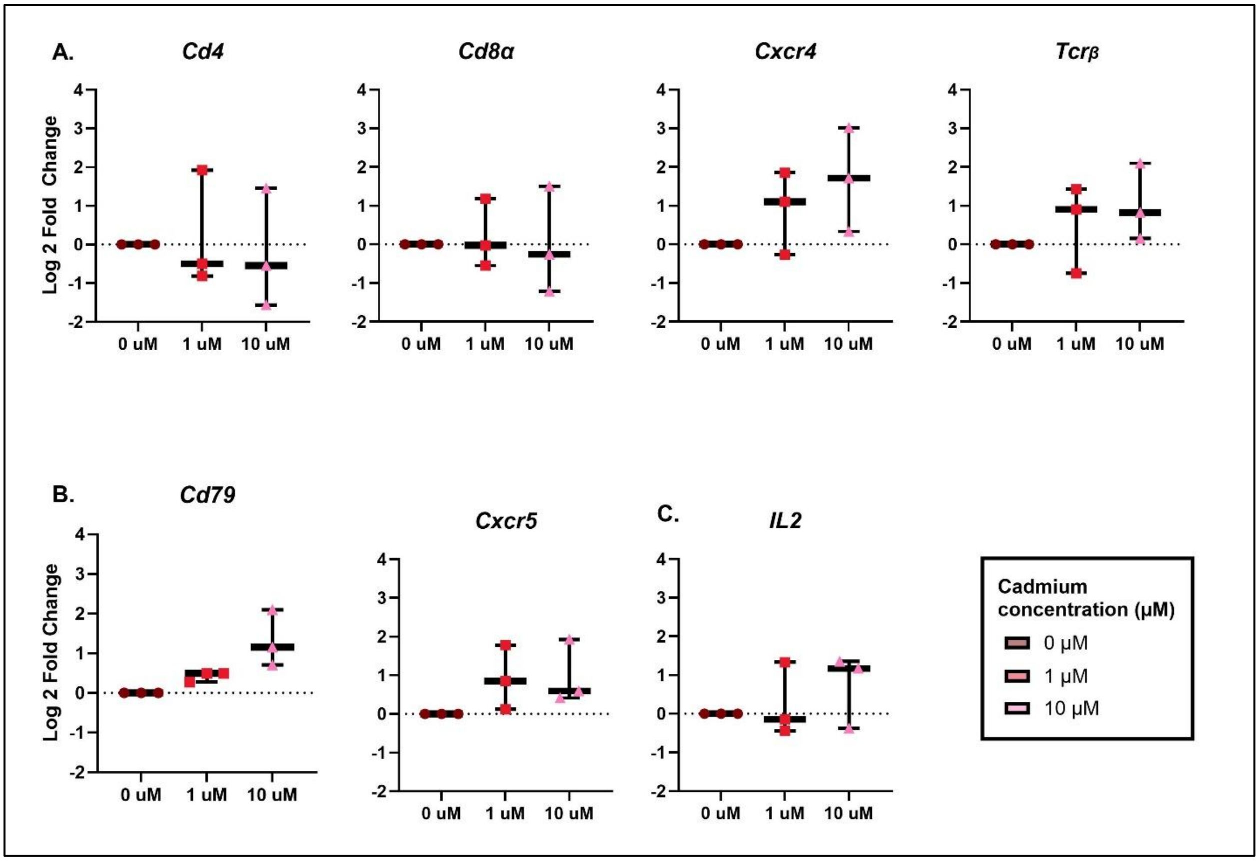


**Supplementary Figure 2:** Log 2-fold gene expression in $CdCl_2$ treated splenocytes from Jamaican fruit bats infected with BANAL 52-Coronavirus (B52-CoV). Panel A) T cells, B) B cells, and C) *IL2* cytokine gene transcripts for splenocytes treated with 0 µM (dark pink), 1 µM (salmon), and 10 µM (light pink) $CdCl_2$. The dashed line represents baseline expression for non-Cd treated cells (0 µM $CdCl_2$ treated group). Fold-change was determined by comparing each transcript from the Cd treated cells to the non-Cd treated cells from the same bat. A mixed-effects analysis with the Geisser-Greenhouse correction and Tukey's multiple comparisons test was used to evaluate significant differences in gene expression between cadmium treatment groups.

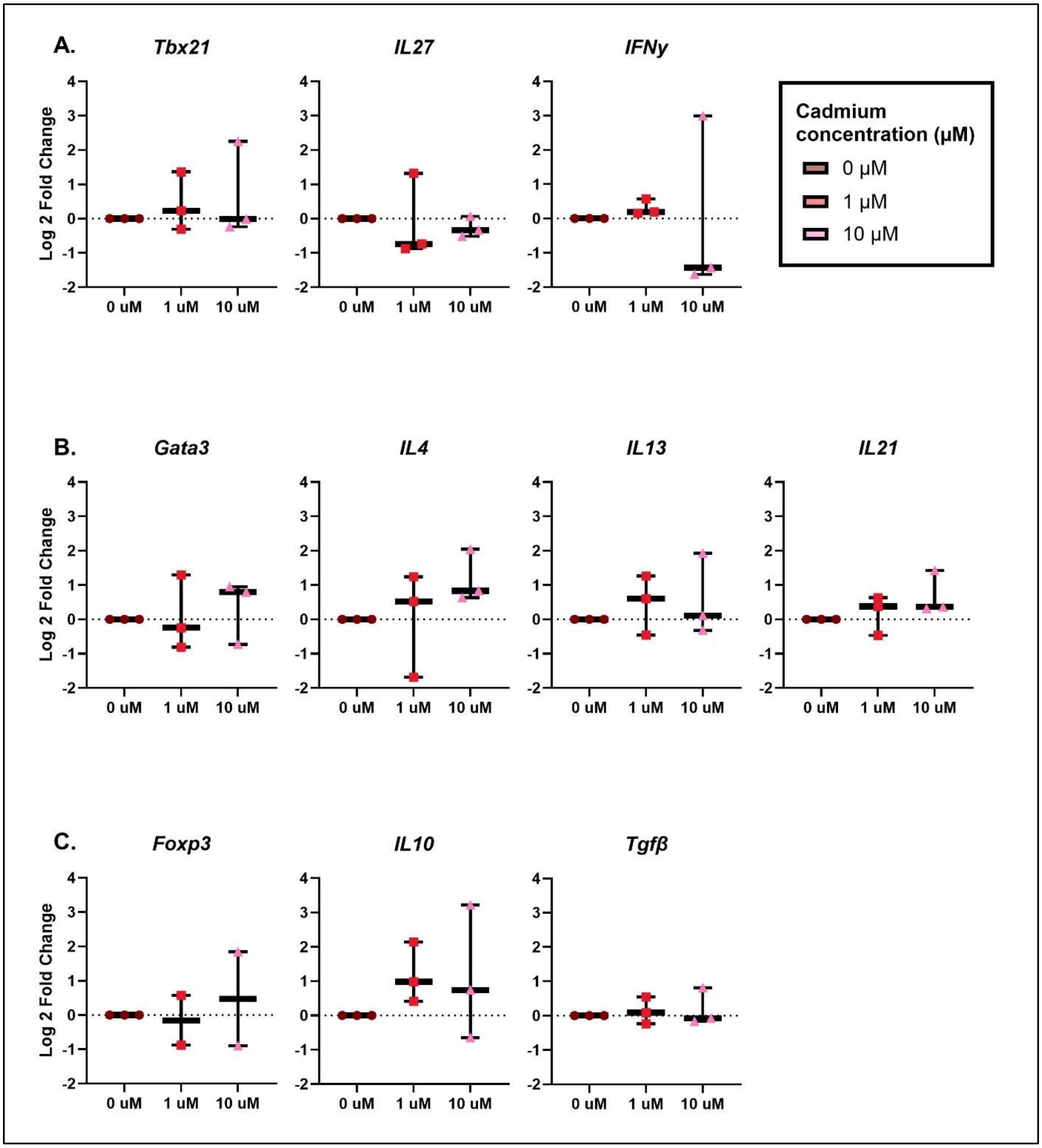


**Supplementary Figure 3:** Log 2-fold gene expression for $CdCl_2$ treated splenocytes from Jamaican fruit bats infected with B52-CoV. Panel A) T helper 1 (Th1), B) Th2, and C) regulatory T cell (Treg) gene transcripts. The dashed line represents baseline expression for non-Cd treated cells (0 µM $CdCl_2$ treated group). Fold-change was determined by comparing each transcript from the Cd treated cells to the non-Cd treated cells from the same bat. A mixed-effects analysis with the Geisser-Greenhouse correction and

Tukey's multiple comparisons test was used to evaluate significant differences in gene expression between cadmium treatment groups.
